# EMS Mutation and SNP Detection in Intracellular *Wolbachia* Genomes

**DOI:** 10.64898/2026.03.29.714874

**Authors:** Gabriel Penunuri, Evan Pepper-Tunick, Jakob McBroome, Russell Corbett-Detig, Shelbi L Russell

## Abstract

Endosymbiotic bacteria such as *Wolbachia* pose significant challenges to genetic and molecular investigation due to their obligate intracellular lifestyle and complex growth requirements.Current understanding of their protein biology relies heavily on functional assignments inferred by homology, which may not reflect the specific roles endosymbiont proteins play within the host. This work addresses the need for robust genetic perturbation by demonstrating the successful application and detection of chemical mutagenesis in the genome of the *w*Mel strain of *Wolbachia* grown within a stably infected *Drosophila melanogaster* JW18 cell line. To accurately detect EMS-induced mutations in a large, unsorted cell culture population, in which mutations remain at very low allele frequency, we implemented an ultra-low error rate sequencing strategy, circle sequencing. This technique enables confident detection of EMS-induced single nucleotide polymorphisms (SNPs) that would be swamped by the inherent error rates of standard next-generation sequencing. Circle sequencing library preparations successfully revealed a clear EMS mutation signal in treated cells, characterized by a significant enrichment of canonical C/G>T/A transitions. Furthermore we present a model explaining observed EMS mutation rates across the genome for different sequence contexts. These findings show that EMS-treatment can successfully leave detectable mutation signals in intracellular genomes, and offer promise for the future development of protocols to make targeted edits in *Wolbachia* genomes.

**Importance:** As the use of intracellular symbionts for bioengineering projects grows, so does the need for foundational protocols for the genetic manipulation of intracellular genomes. Ethyl methanesulfonate (EMS), a chemical mutagen, has been a research tool for initial genomic analysis of gene function in plant and animal systems for decades and represents an established way of generating mutations for future functional testing.

## Introduction

Interest in harnessing intracellular symbiont for large scale biological control continues to grow, particularly as *Wolbachia* demonstrates effectiveness in controlling mosquito-borne diseases and arthropod pest populations ^1–3^. Because of active bioengineering development around obligate intracellular symbionts, particularly *Wolbachia*, accurate functional annotation of endosymbiont genes is critical for progress. Current functional assignments are frequently inferred through homology to genes in free-living organisms, which may not accurately reflect their specific roles within the symbiotic context ^4^.

The absence of robust methodologies for *Wolbachia* genetic perturbation severely constrains the identification of essential genes and the elucidation of specific gene functions ^5^. Intracellular bacteria present significant obstacles to comprehensive investigation; their obligate lifestyle requires complex growth conditions that make genome-wide approaches difficult to scale ^6,7^. Even within stable growth conditions, researchers have found intracellular microbes extremely difficult to genetically manipulate, greatly limiting the scope of experimental interventions ^8^. While impressive advances have been made in obligate intracellular pathogens, such as *Rickettsia* ^9^ and *Chlamydia trachomatis* ^10–12^, progress is slow and expensive. With interest in obligate intracellular symbionts such as *Wolbachia* continuing to increase, the development of a high throughput, genome-wide random mutagenesis methodology would be useful to establishing foundational functional genetic research.

Chemical mutagenesis with ethyl methanesulfonate (EMS) has been a foundational tool in genetics for over a century, enabling systematic functional genetic discoveries across a wide range of plant and animal systems^13^. In contrast, the application of EMS to obligate intracellular bacteria has been limited, with only a small number of studies reporting genome-wide detection of EMS-induced single nucleotide polymorphisms (SNPs)^5,11,12,14^. A primary barrier has been that EMS induces mutations at frequencies three to four orders of magnitude lower than the intrinsic error rates of standard next-generation sequencing platforms^15–17^. This disparity between mutation frequency and technical detectability has prevented the use of EMS in intracellular microbes, despite its proven value for functional genetic analysis in nearly all other biological contexts. Overcoming this limitation requires sequencing approaches capable of substantially suppressing technical error in order to reveal rare, chemically induced mutations.

Here, we overcame these technical barriers by combining chemical mutagenesis with high-fidelity sequencing in a tractable intracellular bacterial system. We leverage a *Drosophila melanogaster* cell line stably infected with the *Wolbachia w*Mel strain, which permits controlled chemical perturbation of the endosymbiont genome. To enable reliable detection of rare EMS-induced variants, we implement circle sequencing^18,19^, a low-error rate library preparation strategy that generates consensus sequences from repeated reads of individual DNA molecules, thereby reducing sequencing artifacts well below standard Illumina error rates. Using this approach, we demonstrate clear detection of EMS mutagenesis signals in treated cell cultures, including enrichment of canonical EMS-associated substitutions and non-random sequence context biases in mutation positions. By demonstrating the first intentional mutagenesis and high-confidence variant detection in *Wolbachia*, this approach establishes a critical prerequisite for future targeted genome editing in these endosymbionts.

## Materials and Methods

### Cell Lines

We maintained *Wolbachia w*Mel-infected immortalized JW18 ^20^ *Drosophila melanogaster* cell lines in 4 mL of media composed of Sheilds and Sang M3 Medium (Sigma S3652) and 10% v/v Fetal Bovine Serum (Gibco A3160502) in plug-seal T25 flasks (Corning 430639), incubated at 25-26°C.

Cell cultures were maintained on a seven-day schedule, splitting at a 1:2 ratio for *w*Mel-infected lines and 1:6 to 1:4 ratio for uninfected lines. For cell passaging, we removed old media, added fresh media, and then dislodged adherent cells by scraping the flask surface with a sterile bent Pasteur pipette (Fisher, 1367820D). Resuspended cells were transferred at the appropriate ratio to new T25 flasks, and the volume adjusted to 4 mL with fresh media. Infection states were monitored with fluorescent *in situ* hybridization against *w*Mel 16S ribosomal RNA and whole genome sequencing (as in ^21^).

### EMS Treatment

We first quantified the *w*Mel-infected JW18 cell concentration in culture with a hemocytometer and mixed media with cells with fresh media in T25 flasks for a final concentration of 1.69e6 cells/mL. These cells were incubated at 26°C for 1 day prior to treatment.

To create EMS-treated and control samples, we added either 20 µl of fresh 0.5% v/v EMS (Sigma M0880) or mock-treatment media, respectively, to 3980 µl of Shields and Sang M3 media supplemented with 10% FBS and sterilized through a 0.2 um syringe filter. We then removed and replaced the old media from previously quantified cell cultures with either EMS or mock media. Flasks were then incubated at 25-26°C for 3, 5, or 7 days.

### Library Prep and Sequencing

We obtained cell pellets from the EMS-treated and controlled cell cultures by scraping the flasks, pipetting 1 mL into 1.5 mL epi tubes, and pelleting the cells by centrifugation at 10,000 × g for 3 minutes at 4°C.

We amplified genomic DNA templates by rolling circle amplification (RCA) using the procedure described in the initial circle sequencing publication by Lou et al. 2013^18^, with several adjustments. DNA purification steps were done with a SPRI magnetic bead protocol instead of the QIAGEN MinElute clean up kit. We sheared gDNA resuspended in TE buffer over two rounds of five minutes with a spin down between rounds. We set the duty cycle to 30%, peak power to 60W, and 1000 cycles per burst. Phosphorylation was done with the T4 PNK as described in the supplemental methods of Lou et al. ^18^. The DNA was then denatured and flash frozen according to the Lou et al. protocol. For the circularization reactions we used the same master mix except we added no more than 3.5 pmol of ssDNA instead of 5 pmol. Linear DNA remaining in the reaction mix was digested according to the Lou et al. protocol. For the rolling circle amplification (RCA) reactions we skipped the random primer annealing reaction, and instead used 2.5 μl of 10 μM random hexamer primers in the RCA reaction mix. We used 2.5 μl (instead of 1 μl) of 10 mM dNTP and did not include the inorganic pyrophosphatase. After performing RCA, we treated about 20-30 ng of RCA DNA products with 1 μl of DNA Exonuclease I (NEB M0293S) and 0.5 μl of DNA Exonuclease III (NEB M0206S) for 1 hour at 37ºC to digest linear fragments. We then purified samples using SPRI magnetic beads and Qubit quantified with the dsDNA HS Assay Kit (Thermo Fisher Q32851).

We prepared libraries from the RCA products by mixing 10 ng of RCA products with 2 μl charged Tn5 transposase (Tn5 enzyme was expressed and purified in-house) in 12 μl of molecular grade water and 4 μl of 5x TAPS-PEG 8000 for 8 minutes at 55ºC ^22^. The reaction was halted by adding 0.2% SDS and incubated at room temperature for 7 minutes. We set up PCR reactions using the KAPA Biosystems HiFi Polymerase Kit (Roche) and amplified for 12 cycles using uniquely indexed i5 and i7 primers. We ran these PCR products on a Thermo Fisher 2% E-Gel EX Agarose Gel (G401002) and cut between∼400-700 bp. The gel was purified with the NEB Monarch DNA Gel Extraction Kit (NEB #T1020) and quantified using the dsDNA HS Assay Qubit Kit and the Agilent TapeStation D1000 tapes (5067-5582). We prepared two pools of circle sequencing Tn5 libraries: one pool containing 14 purified and indexed libraries from 5-day samples, and a second pool containing 14 purified and indexed libraries from 3 and 7-day samples. We sequenced both pools using 150-bp paired-end reads across two Illumina HiSeq 4000 lanes at Fulgent Genetics.

### Circle Sequencing Processing

We processed the demultiplexed circle sequencing short read data using a custom snakemake ^23^ pipeline (https://github.com/jmcbroome/circleseq). We aligned paired-end sequencing FASTQ reads to the *w*Mel *Wolbachia* reference genome (NC_002978.6) using BWA-MEM^24^, and output the alignments as SAM files. The aligned sequences were divided into repeat subsequences based on their mapping, with trimmed or unaligned repeat sequences separated from the primary alignment with custom python scripts. We selected and extracted reference genome regions with BEDTools getfasta ^26^ based on the genomic coordinates of the primary alignments generated from the BWA mapping step. Groups of subsequences were then realigned to their mapping region using an optimized striped Smith-Waterman alignment provided by scikit-bio^26^. For each group, we constructed a consensus sequence from the realigned subsequences and defined the quality score as the number of times the base was observed. Any base with a mismatch in a subsequence realignment was replaced with an N. In the final step, the consensus alignments were piled up with SAMtools mpileup^27^, after filtering positions with less than two subreads present.

This approach, although sharing the same core idea of implementing error-suppression with consensus construction, differs slightly from the original methods published by Lou *et al*. ^18^. We have opted to use reference-anchored repeat detection through BWA-MEM alignment instead of self-alignment between paired-end reads. Our approach also differs by how the consensus is built. Originally published methods did so from explicit tandem copies where each repeat corresponds to a single pass around the circularized read leading to repeat boundaries that are regular and periodic. Our approach builds the consensus from subsequences extracted after mapping resulting in length variation that is more tolerant of irregular repeat structure which can better reflect artifacts of Illumina library preparation. We also implement a two pass alignment strategy that allows for greater specificity and scalability for genome-wide analysis through performing a local Smith-Waterman realignment after the initial global BWA-MEM alignment.

This step further builds analysis tolerance for truncated repeats, uneven coverage and sequencing dropouts that would all be expected from Illumina short read sequencing. It is also during this secondary alignment step that we increase the specificity of variant detection and lower the false-positive rate. We are able to track any disagreement across the subreads and mark these potential PCR-induced errors or alignment artifacts by calling the bases as N removing them from downstream analysis.

### Mutation Rate Estimation Per-Sample Rates

We collected mutation counts from mpileup files with a custom python script (https://github.com/gabepen/ems_effect). We loaded site-level mutation counts from these mpileup files. Sites were restricted to G/C positions. For C sites, only C→T mutations were counted; for G sites, only G→A mutations. We constructed an exclusion mask representing genomic positions where mutated sites appeared in more than one control sample. We used this mask to exclude these positions from all downstream analysis. We extracted sequence contexts (3mers, 5mers and 7mers) from the reference genome, and G-centered contexts were reverse-complemented to C-centered for canonicalization. We used a finite-sites model to estimate an initial per-sample mutation rate for mutation spectra visualization:

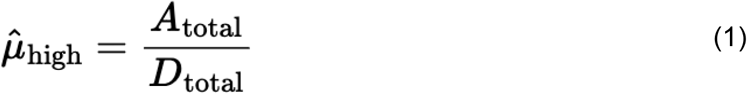

Where A_total_ is the count of alternative alleles across all positions and D_total_ is the total sequencing depth across all sites.

We modeled per-sample mutation rates using generalized linear models (GLMs) with log(depth) as an offset starting with a background subtraction:

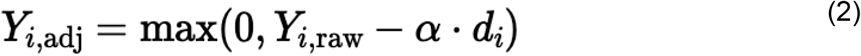

These adjusted counts are then used in a GLM:

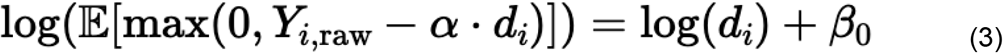

Where:

*Y*_*i,raw*_ = raw mutation count at a site *i*

*Y*_*i,adj*_ = alpha-adjusted mutation count at a site *i*

*d*_*i*_ = sequencing depth at site *i*

*α* = background false positive rate (estimated from controls)

*β*_0_ = intercept parameter, estimated per sample

We then estimated rates as:

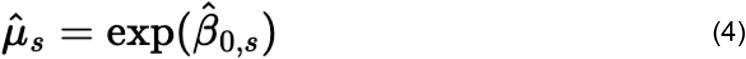

### Mutation Rate Estimation Across Genomic Categories

We estimated mutation rates for intergenic, synonymous, and non-synonomous rates with a negative binomial GLM. We opted to use a negative binomial distribution to account for the overdispersion and low mutation counts in the finite-sites dataset. We fit this model with treatment and category covariates, including interactions:

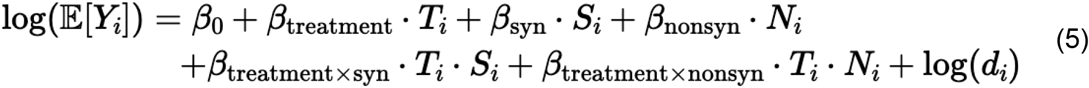

We modeled these same genomic categories with an alternative approach to capture the absolute instead of the proportional effect of treatment on category rates starting with a control only model to establish a baseline:

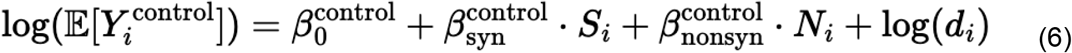

Next we calculate the expected mutation excess with treatment:

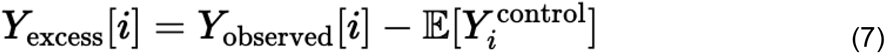

We then fit two GLMs using Gaussian family with an identity link, the first a model for constant absolute effect:

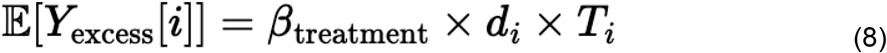

And second an interaction model:

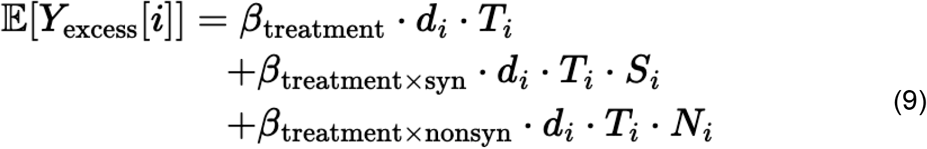

We also estimated category rates for treatment time with a negative binomial GLM, where no-label is the 5 day reference:

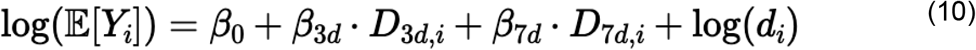

### Mutation Sequence Context Analysis

We modeled per-site EMS mutation counts using generalized linear models (GLMs) with sequence-context features.

Site-level mutation counts were loaded from mpileup files. Sites were restricted to G/C positions. For C sites, only C→T mutations were counted; for G sites, only G→A mutations. Sites in an exclusion mask (those appearing in multiple control samples) were excluded. Sequence contexts (3mers, 5mers and 7mers) were extracted from the reference genome, and G-centered contexts were reverse-complemented to C-centered for canonicalization.

We used Negative Binomial GLMs to model the sequence context of EMS mutations. The per-site EMS mutation count served as the response variable, and log(depth) was included as an offset term. The model evaluated the effect of the experimental conditions by including both the treated and control groups as primary covariates.

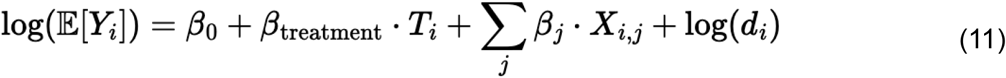

Where:

*Y*_*i*_ = raw mutation count at a site *i*

*T*_*i*_ *=* treatment indicator

*X*_*i,j*_ = sequence context features (vary by model type)

*d*_*i*_ = sequencing depth at site *i*

*β*_treatment_ = treatment effect on log scale

*β*_j_ = sequence context coefficients

*β*_0_ = intercept parameter, baseline log rate for control sites

To account for sequence context dependencies and to determine what context was the best predictor of mutation bias, we implemented five distinct encoding schemes:

1. Positional: We encoded sequence context using 12 binary features representing the base identity (T, G, or C) at each of the 4 varying positions in the canonicalized 5-mer, with A as the reference category for each position. The center position is always C, so 3 features are constant; 12 features vary across sites. Coefficients represent log-enrichment relative to A at each position.
2. 3-mer: We encoded context using 16 binary features representing all possible C centered 3-mers spanning positions 1 through 3.
3. 5-mer: We encoded context using a one-hot encoding approach, resulting in 256 binary features.
4. Positional-3-mer: We encoded context using 32 binary features, representing the left and right 3-mers covering the -2, -1, 0 and 0, 1, 2 positions respectively. This model also evaluates C centered mutations only resulting in fixed ‘0’ positions in both left and right 3-mers.
5. 7-mer: We encoded context using a one-hot encoding approach, with a theoretical maximum of 4,096. To manage memory usage, the model required the input contexts to be split into separate groups deterministically by first sorting unique 7-mers. Model fitting was performed independently for each split, and the resultant metrics were subsequently aggregated to obtain the final summary statistics.

After fitting, we predicted mutation counts for each site under control and treated conditions. We converted predicted counts to per-base mutation rates by dividing by depth, then aggregated observed and predicted rates by the relevant sequence-context category for the model using depth‐weighted means. We evaluated fit by comparing observed versus predicted aggregated rates using scatter plots and correlation metrics.

### Cross-Validation Evaluation (Out-of-sample)

We used 5-fold cross-validation to measure out-of-sample predictive performance. We split sites into five folds with sklearn.model_selection.KFold using shuffling (random_state=42). For each fold, we trained each model on four folds and evaluated it on the held-out fold, so every site served as test data exactly once. For each held-out fold, we computed mean squared error (MSE), mean absolute error (MAE), Pearson correlation (r), and a negative-binomial deviance statistic. We summarized performance across folds by reporting the mean (and, when available, the standard deviation) of each metric.

For the 7-mer split model, we controlled memory usage by fitting the model in groups of 7-mer categories. We deterministically partitioned unique 7-mers into groups using a round-robin assignment over sorted 7-mer categories. Within each cross-validation fold, we fit one GLM per group using only training sites in that group. We generated held-out predictions by applying the corresponding group model to held-out sites in that group (each site received a prediction from exactly one split model), and we computed cross-validation metrics from the concatenated held-out predictions across groups.

### Prediction Accuracy Metrics (Evaluation dataset; in-sample in the standard pipeline)

We also evaluated prediction accuracy on the evaluation dataset by comparing observed mutation counts (y) to model-predicted expected counts (mu) while incorporating the log(depth) offset. For the 7-mer split model, we produced predictions by routing each site to the split model corresponding to its 7-mer category, and we evaluated performance only on sites with a valid 7-mer context.

We computed MSE, MAE, RMSE, and MAPE; Pearson correlation (r) and Spearman correlation (rho) with p-values; R-squared and adjusted R-squared; a deviance statistic; MSLE and RMSLE; and median-threshold classification metrics (accuracy, precision, recall, and F1). We also reported the number of evaluated sites and the mean observed and predicted counts.

### Genome-wide EMS mutation enrichment

We analyzed local genomic variation independent of genomic categories by dividing the genome into fixed-size, non-overallaping 5-mer windows and testing if the observed rates deviated from the 5-mer model for EMS mutation context and rate.

For each mutated site *i* in window *W*_*k*_ we extract the 5-mer context and predict the expected mutation rate using the pre-fitted 5-mer model. We then fit a site-level GLM with a 5-mer offset and a treatment covariate:

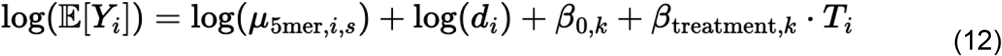

Where:

*Y*_*i*_ = observed mutation count at a site *i*

*T*_*i*_ *=* treatment indicator

*µ*_5mer,i,s_ = pre-computed expected rate from the 5-mer model

*d*_*i*_ = sequencing depth at site *i*

*β*_treatment,k_ = window-specific treatment effect

*β*_0,k_ = window-specific intercept

### Genome-wide EMS mutation enrichment in the context of transcription

We tested the correlation between EMS mutation enrichment and rates of transcription by adapting the fixed-size window models above to encompass genic regions and performed a regression analysis between the predicted window mutation rates and TPM values for each corresponding gene. To obtain TPM values for the wMel genome, we downloaded the wMel-infected JW18 cell RNAseq dataset produced by Jacobs *et al*. (2025) (PRJNA1240446) and regenerated pseudoalignments according to their methods^28^. We used ordinary least squares (OLS) linear regression fitting controls and treated samples separately.

## Results

### EMS Mutation Signal

Using high-fidelity circle sequencing^18^, we identified an elevated mutation rate in the EMS-treated *w*Mel genome cultured in *Drosophila* JW18 cells, relative to control treatments. Following Illumina sequencing and demultiplexing of the 18 EMS-treated and 10 control samples, we obtained 4,691,282,072 total reads estimated to be derived from 619,355,451 rolling circle amplification (RCA) products amounting to 2,755,257,101 total read copies (Table S1, S2). When we compared EMS-treated and control consensus circle sequencing reads to the *Wolbachia w*Mel reference genome, GCF_000008025.1, treated samples showed a significant enrichment of the canonical EMS-induced transitions C>T and G>A (p<0.001, Mann-Whitney U test, Figure 1A). These mutations affected the entire genome (Figure S1, Table S3). We also noted changes to the physical characteristics of the EMS-treated cell lines when compared to the control lines (Figure 1B, 1C). Treated cells exhibited morphological signs of stress including slowed growth, small cell size, condensed cytoplasm, and disrupted membranes.

**Figure 1.**
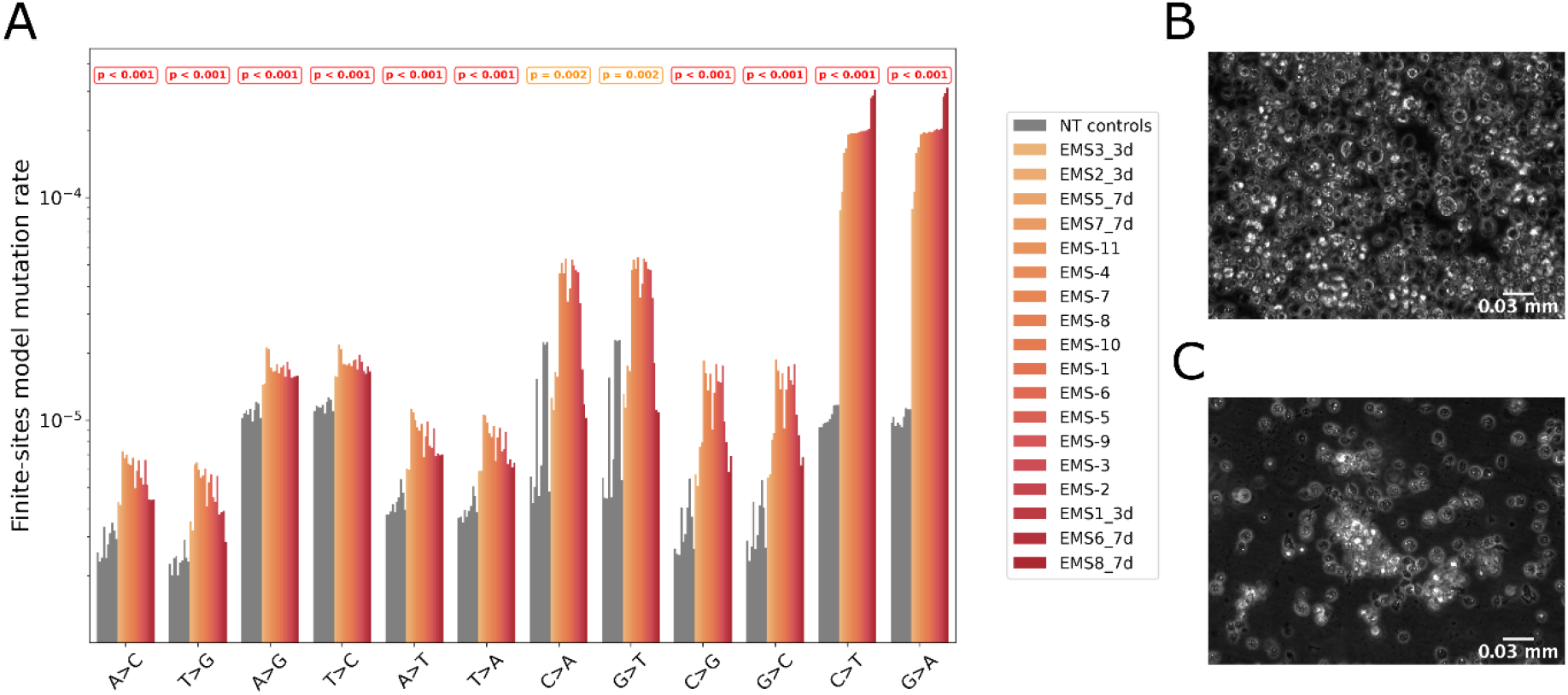
Treating *w*Mel-infected JW18 cells with EMS induces C>T and G>A mutations in the *w*Mel genome. Representative tissue culture micrographs of *w*Mel-infected JW18 cells **A)** after the control treatment and **B)** after EMS treatment. **C)** EMS mutation spectra demonstrating a large difference between control (grey) and treatment (orange-red gradient by experiment) at the canonical EMS transitions types of C>T and G>A.

We represented the mutation spectra as a depth-normalized proportion of sites mutated for each transition type using the finite-sites model, indicating that the EMS mutation signal was only clearly detectable through low error rate sequencing methods (Figure 1C). In previously published mutation rates, EMS has been shown to induce per-site mutation rates of ∼1 × 10^−6^ to ∼1 × 10^−2^ across different organisms ^15,16^ while the minimum medium error rate on an Illumina platform is around 5 × 10^−3^ per-site ^29^.

### EMS Mutation Rates

We estimated mutation rates across all samples using site-level GLMs, obtaining estimates of per-sample rates ranged from 2.23 × 10^−5^ to 3.17 × 10^−5^ mutations per base pair for controls (mean: 2.71 × 10^−5^ mutations/bp) and 9.44 × 10^−5^ to 3.24 × 10^−4^ for treated samples (mean: 2.05 × 10^−4^ mutations/bp). Treated samples exhibited significantly elevated mutation rates with average increase of 4.73-fold (95% CI: 4.65 - 4.80) compared to controls (p <0.001, negative binomial GLM with treatment covariate).

The treatment durations used in the EMS dosage protocol had a proportionate effect on mutation rate when we modelled the interaction between treatment duration and mutation rate using a negative binomial GLM with days of treatment as a covariate. We grouped control and treated samples as 5-day (treated=11, control=3), 3-day (treated=3, control=3), and 7-day (treated=4, control=4). All pairwise comparisons between groups were significant with the group-level estimates showing an increase in mutation rates as treatment time increased (Figure 2C). However, we note substantial inter-sample variance over each treatment duration category.

**Figure 2.**
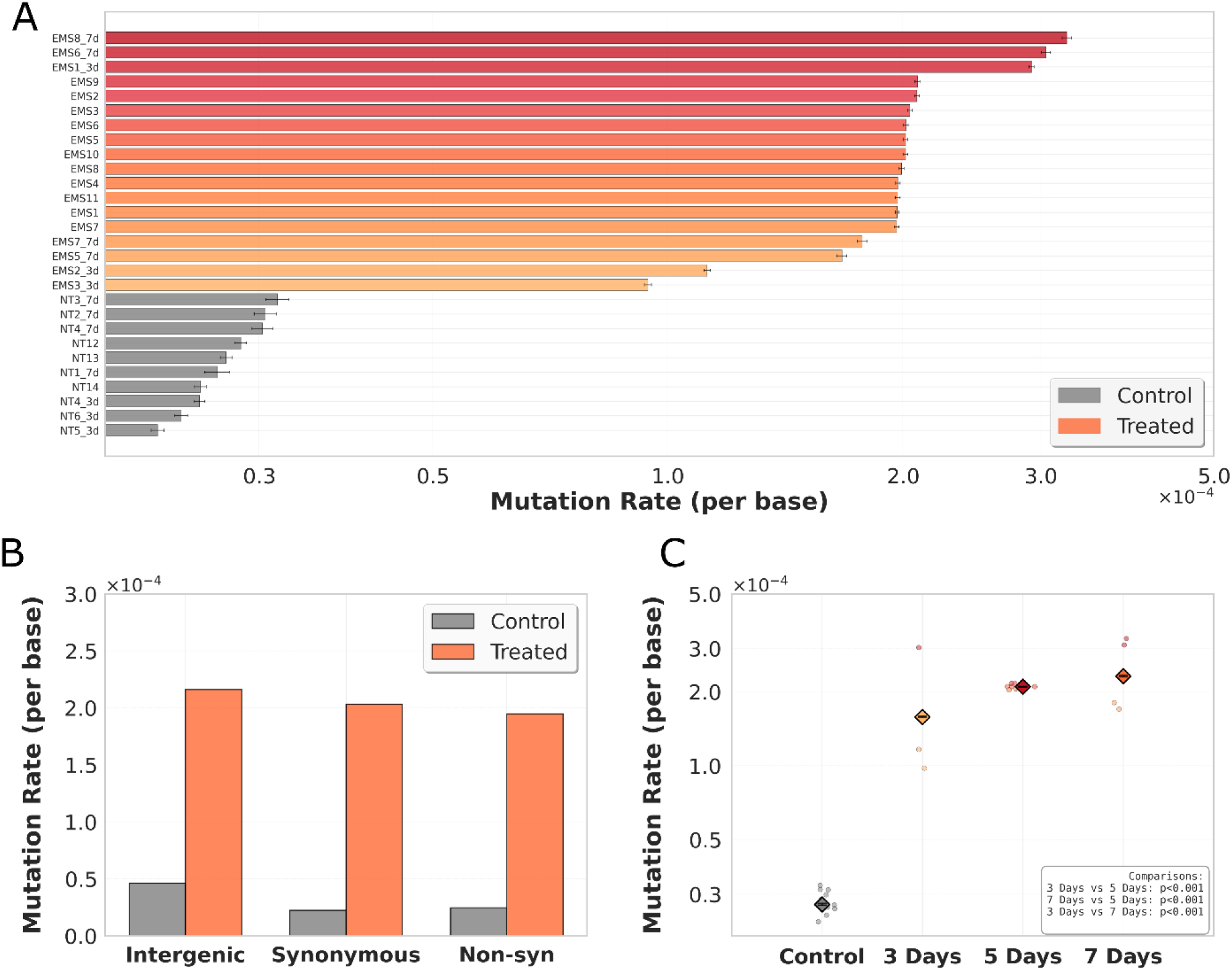
EMS treatment duration was positively associated with mutation rate, whereas functional aspects of the wMel genome exhibited no association. **A)** Per-sample EMS mutation rates for treated and control samples as estimated by fitting a negative binomial GLM. Treated samples exhibit∼5-fold higher mutation rates than control samples. **B)** Intergenic versus coding mutation rate analysis with treated rates normalized by 5-mer content in each region category. Similar absolute rate differences among coding categories suggests that EMS treatment added the same number of mutations per base pair across genomic categories. **C)** Treatment time had a significant impact on mutation rate when treatment time was modeled as a covariate of mutation rate, with longer treatment times correlating with higher mutation rates.

We determined that the functional sequence category has a negligible differential effect on EMS mutation incidence by comparing mutation rates across three different functional coding categories: intergenic, synonymous and non-synonomous. We calculated mutation rates by incorporating a baseline mutation rate for each genomic category using the 5-mer model described in the mutation sequence context analysis portion of the methods. This accounted for any significant divergence in sequence content between coding and intergenic regions. We estimated control mutation rates for these three categories to be 4.63 × 10^−5^, 2.24 × 10^−5^ and 2.26 × 10^−5^ mutations per base, respectively. We estimated treated rates to be 2.16 × 10^−4^ mutations per base pair for intergenic sites, 2.03 × 10^−4^ mutations per base pair for synonymous sites and 1.95 × 10^−4^ mutations per base for non-synonomous sites (Figure 2B). Although these rates represent large differences between categories in the proportional changes between the control and treated samples they amount to similar absolute effect, as quantified by the number of mutated bases added by EMS to the background control rates. For the same three categories, the absolute EMS effect, or mutations per base added by treatment, were 1.16 × 10^−4^, 1.73 × 10^−4^, and 1.62 × 10^−4^. While the constant effect and interaction models produce statistically significant effect differences (p < 10^−154^, likelihood ratio test), they only explain about∼5% of the variation. This indicates that the rate differences between intergenic and genic regions are biologically insignificant.

We found no evidence to suggest that EMS is preferentially mutagenizing regions of the *w*Mel genome that are open for transcription. We calculated EMS mutation rates within 2500 base pair non-overlapping windows across the genome using our k-mer aware genomic window rate model (Eq. 12) to identify biologically meaningful hot or cold spots. A binomial test of window clusters, comparing mutation rates to the genome-wide mean, identified no stretches of EMS hot or cold spots beyond a single window. We also calculated EMS mutation rates for each gene and performed a regression analysis with *w*Mel *Wolbachia* gene expression data collected by Jacobs et al. (2025)^28^. The expression data were weakly correlated with EMS mutation rate (R^2^ = 0.025, Figure S2). This combined with the random spatial distribution of significantly elevated windows (with no detected clustering) suggests that mutation rate variation in this system is influenced by unmodeled effects. While the treatment induces a global increase in mutation rates, the local variation appears to be largely random, with no evidence for transcription-associated mutagenesis.

### EMS Mutation Sequence Context Bias

We selected the 5mer model for determining sequence bias in EMS mutations because it provided the best balance between model fit and generalizability across multiple performance metrics (Table S4). The 7mer model achieved the lowest AIC among all evaluated models with a ΔAIC to the next lowest, the 5mer model, of∼18,000 (Table S4). However, the 7mer model also reported the highest overall BIC indicating the capturing of a significant amount of noise. The 5mer model reported the lowest BIC with a ΔBIC of∼5000 to the second lowest, the positional-3mer. Thus our prediction accuracy and cross-validation tests supported the 5mer model over the 7mer, 3mer, and positional models (Figure S3). For prediction accuracy the 5mer model had the lowest RMSE (0.546), and highest pearson R (0.348) with the positional-3mer model performing second best (RMSE = 0.299, pearson R = 0.343) and the 7mer third (RMSE = 0.300, pearson R = 0.342); while simpler models (positional, 3mer) showed reduced predictive accuracy. Cross-validation (CV) metrics further suggested although the 7mer model was adept at producing rankings, overfitting likely hindered its calibration. The 7mer model had the highest CV-MSE (R=0.319) and lowest CV-pearson (R=0.339). The 5mer model performed the best in these metrics with R=0.298 and 0.367, respectively.

All models performed poorly at predicting EMS mutations from sequence context alone, indicating that additional factors contribute to EMS mutation bias. Nevertheless, our fitted 5mer model identified significant enrichment of certain sequence contexts at EMS-mutated sites (Figure 3A, Table S5). AT dinucleotides were enriched at the -2 and -1 positions directly upstream of EMS-mutated bases (Figure 3C), while dinucleotides containing A or T were depleted at the downstream +1 and +2 positions where C or G dinucleotides were enriched. Our results suggest that although C and G bases are enriched near EMS-induced SNPs, this enrichment occurs predominantly downstream rather than uniformly surrounding the mutated base.

**Figure 3.**
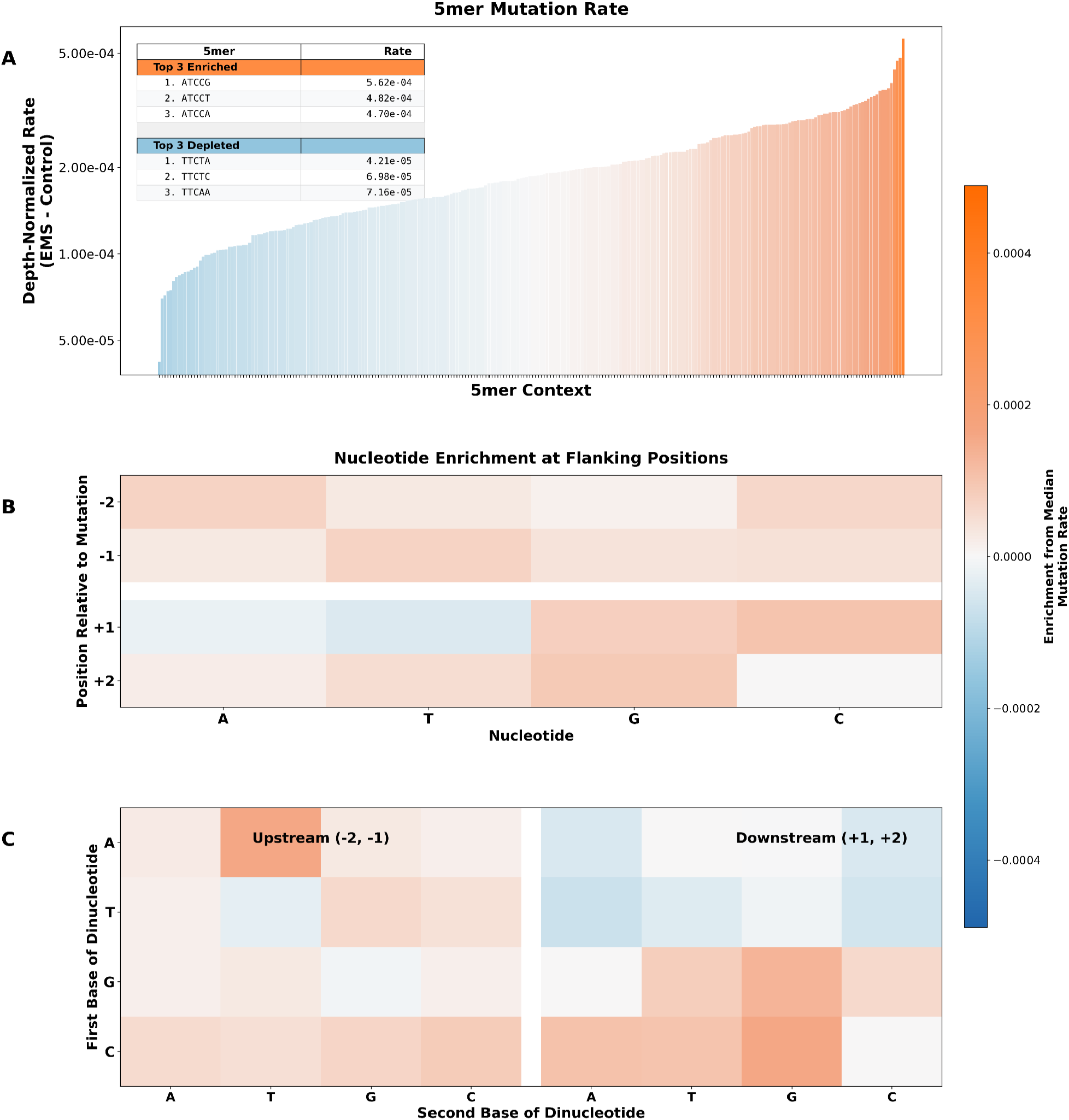
EMS mutations in the wMel genome exhibit biased sequence context. **A)** Mutation rates of all possible 5mer sequences. Strands are collapsed, such that all canonical EMS mutations are treated as C-centered. The top three enriched and depleted sequences are shown in a table with the associated mutation rate. **B)** Single nucleotide enrichment at each of the four positions flanking a canonical EMS mutation site. **C)** Dinucleotide pair enrichment at the upstream and downstream positions flanking an EMS mutation site.

## Discussion

Researchers have made significant advances in the tractable genetic perturbation of obligate intracellular bacteria ^5^. However genetic perturbation of diverse intracellular bacterial symbionts remains an extended goal of researchers looking to better understand and harness their unique biological niches. Obligate intracellular symbionts like *Wolbachia* are already being used in large scale biological control projects seeking to improve both agricultural and human health ^1,3^ yet approaches to modify their genomes are lacking. Here we validated the use and detection of EMS for random chemical mutagenesis of intracellular *w*Mel *Wolbachia* infecting *Drosophila* cell culture. Using an extremely low error rate sequencing method ^18^, we produced mutation profiles with a nearly 5-fold increase above the background mutation and sequencing error rate.

Together, these results establish EMS mutagenesis as a tractable and quantifiable approach for generating genome-wide genetic variation in intracellular *Wolbachia*, providing a practical foundation for future functional genetic studies in systems where genetic manipulation has historically been inaccessible.

Beyond establishing the feasibility of detecting EMS-induced mutations in an intracellular genome, our results suggest that EMS imposes a largely uniform mutational burden across functional genomic categories. The minimal difference observed between coding and non-coding mutation rates indicates that EMS does not preferentially target or spare protein-coding regions, consistent with broadly stochastic mutation rather than context-specific targeting or repair. This uniformity is further supported by the absence of extended mutational hot or cold spots across the genome, arguing against large-scale regional variation in EMS accessibility. This implies that, with sufficient mutational saturation, EMS mutagenesis is expected to impact essentially all genomic regions in *Wolbachia*, making it well suited for genome-wide functional interrogation.

At a finer scale, the lack of a strong relationship between gene expression and mutation rate suggests that transcription-coupled processes play a limited role in shaping EMS mutational outcomes across the *w*Mel genome. Nevertheless, clear sequence-context preferences remain evident, with a 5-mer model providing the most informative description of EMS mutation bias. The fact that even this model performs modestly underscores that EMS mutagenesis is influenced by factors beyond local sequence alone, likely reflecting interactions with genome organization or DNA repair pathways. Together, these findings highlight both the predictability and the limits of sequence-based models of EMS mutagenesis in intracellular genomes.

Our work proposes a future for genomic screens in *Wolbachia* as well as the potential within other host-cell cultured endosymbionts. This demonstration of high-confidence detection of EMS-induced mutations in *Wolbachia* using ultra-low error rate sequencing allowed us to build on previous models of EMS mutation rate and bias, and it contributes towards future use in larger screens. Refined EMS dosing techniques and methods could produce saturated mutation datasets allowing for the description of essential genes in symbiont genomes, functional gene annotation and further phenotypic based screens.

## Acknowledgments

This work was supported by NIH R00GM135583 and R35GM157189 to SLR as well as T32HG012344 to GP.

## Data Availability

All code used for processing circle sequencing data can be found here: https://github.com/jmcbroome/circleseq

All code used for rate modeling, analysis and data visualization can be found here: https://github.com/gabepen/ems_effect/tree/manuscript-scope

All raw sequencing data files can be found under NCBI Bioproject PRJNA1427293

## Supplemental Materials

**Table S1. Read Counts**. Per-sample initial reads, RCA products and total read copies.

**Table S2. Per-sample Metrics Table S3. Per-gene Metrics**

**Table S4. Model Selection Summary Statistics Table S5. 5-mer Sequence Bias Model Rates**

**Figure S1.**
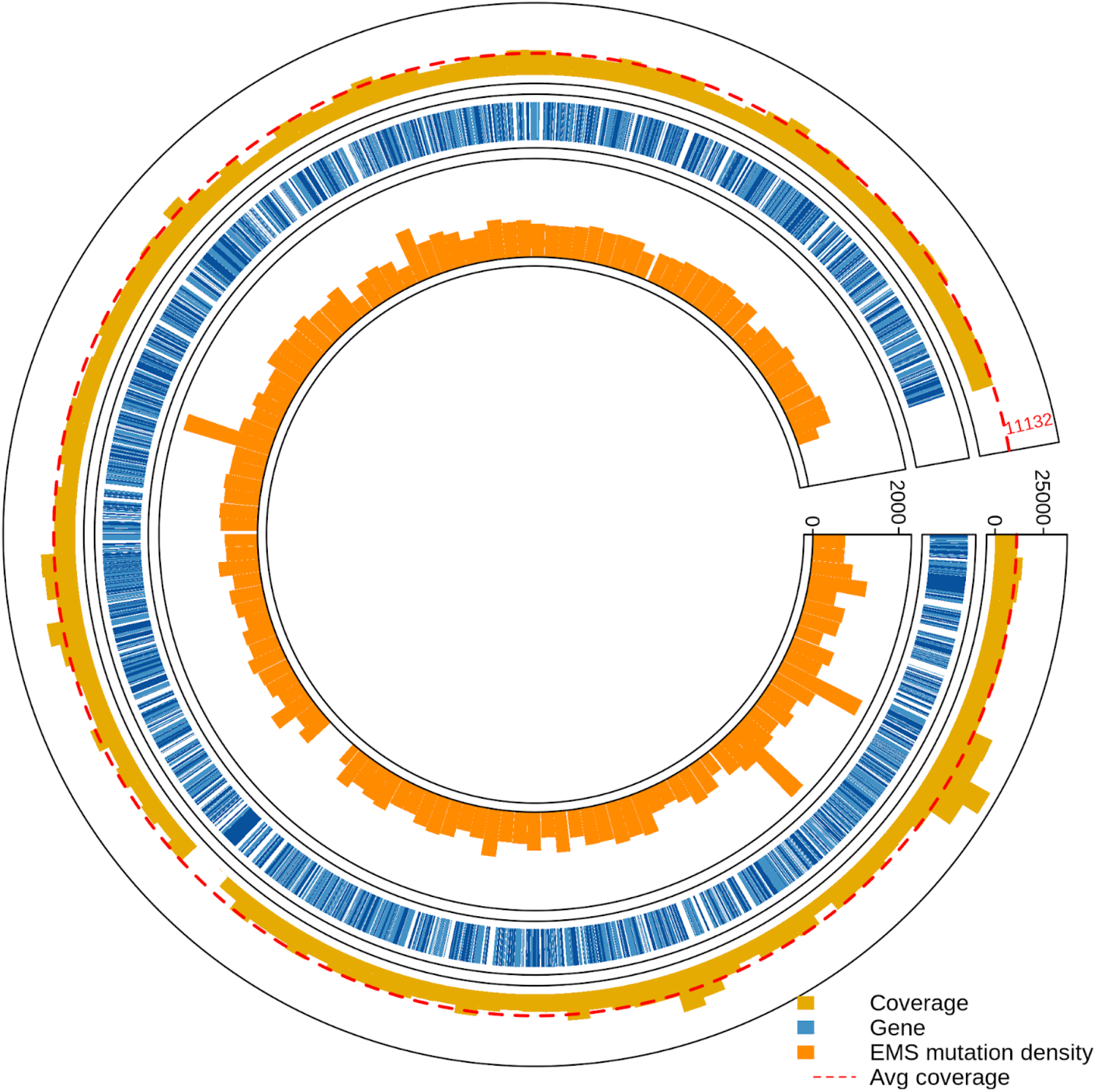
EMS mutation density across the genome. The bars represent 10,000 bp windows. The gap in mutation density coincides with no sequencing coverage. This is due to the drop out of the octomom gene cluster from the JW18 hosted wMel cell line.

**Figure S2.**
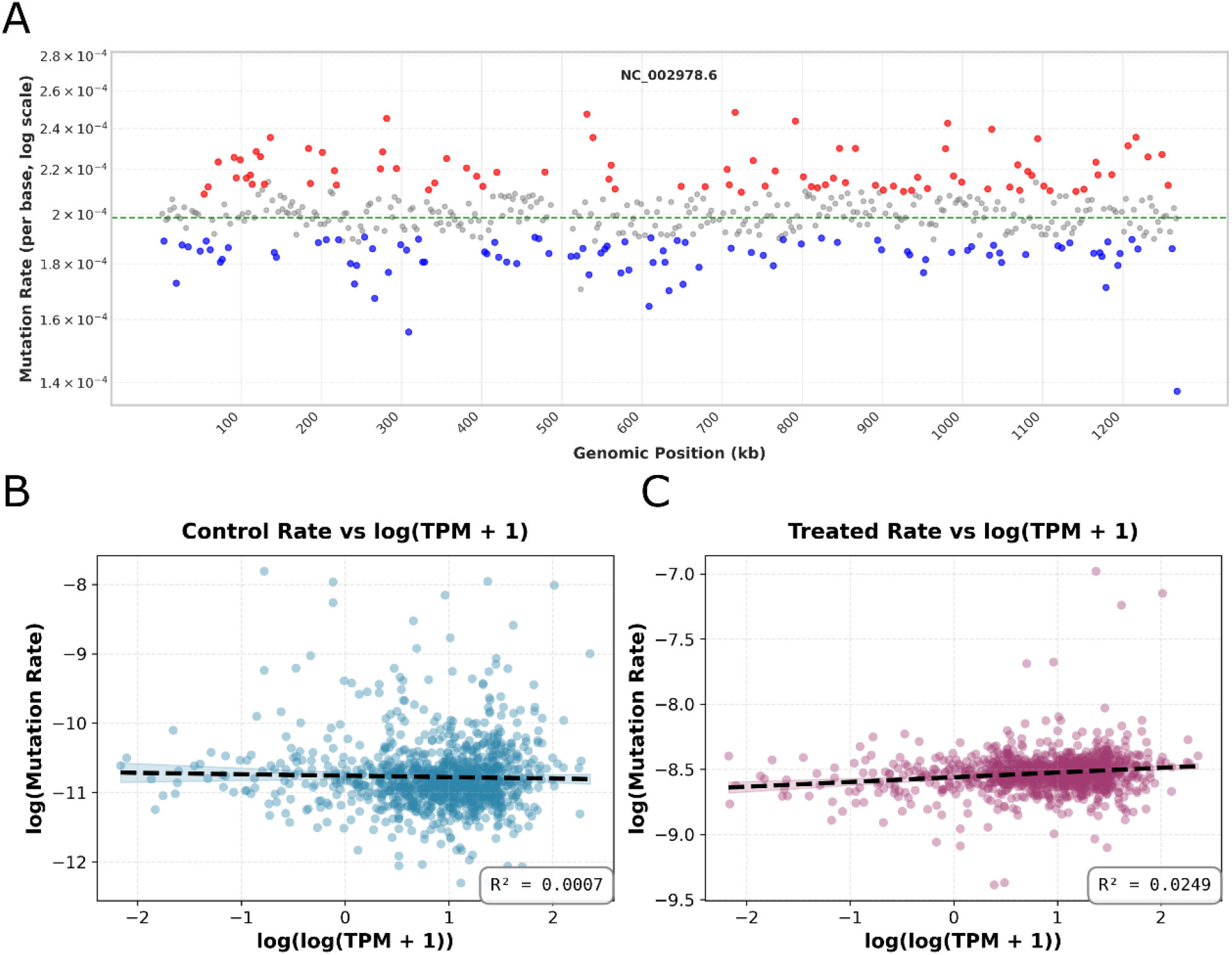
Analysis of EMS Mutation Rates Across Genome and Genes. **A)** Plot of 2500 bp windows that have a significantly higher (red) or lower (blue) mutation rate compared to the overall mean. **B, C)** Linear regression analysis of control and treated rates for genes against their expression levels.

**Figure S3.**
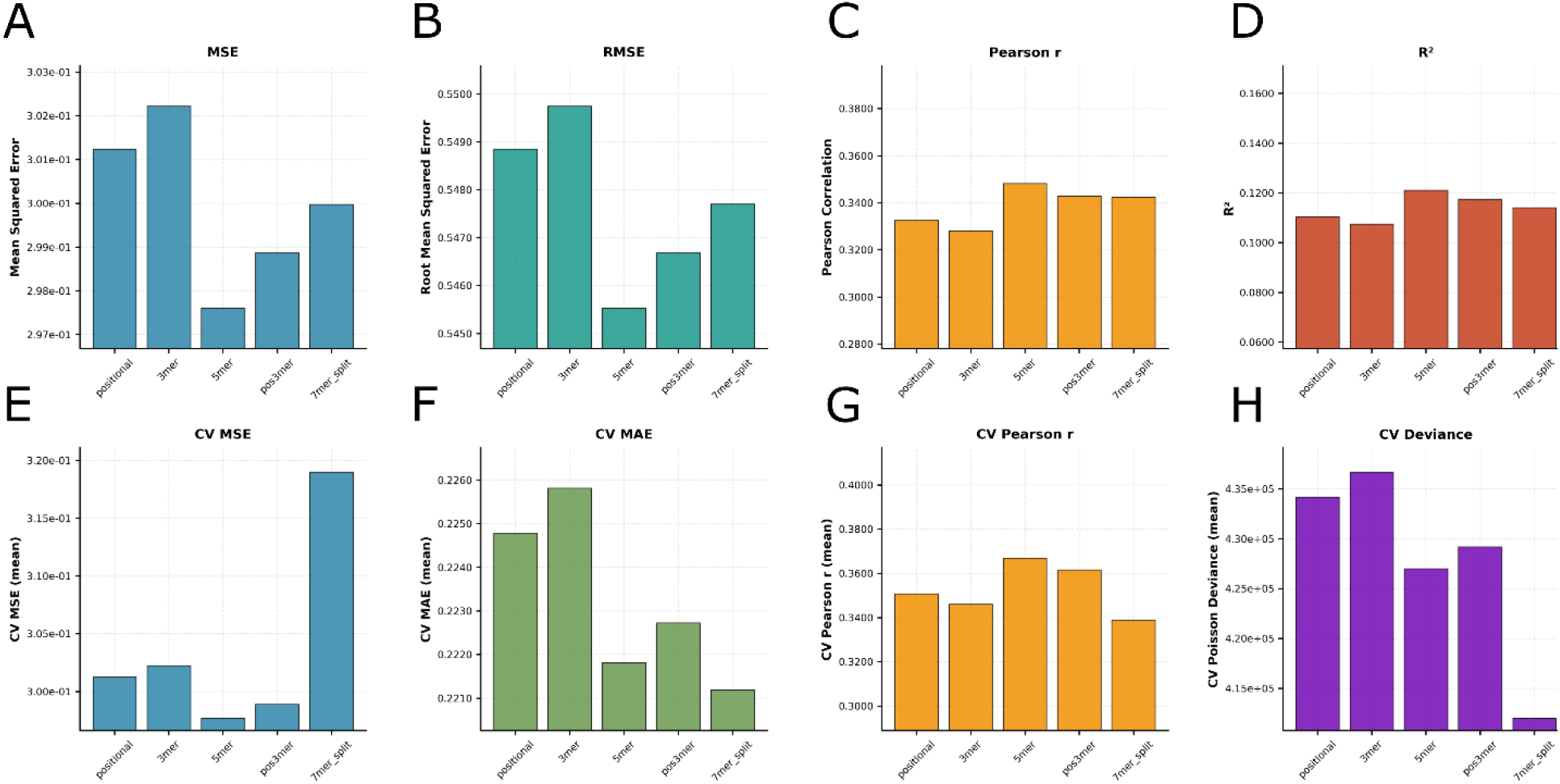
Sequence Bias Model Evaluation. **A-D)** Within sample prediction accuracy metrics for each model type. **E-H)** Out of sample cross-validation metrics for each model type

